# White Matter Microstructure Correlates of General and Specific Second-Order Factors of Psychopathology

**DOI:** 10.1101/459396

**Authors:** Kendra E. Hinton, Benjamin B. Lahey, Victoria Villalta-Gil, Francisco A. C. Meyer, Leah L. Burgess, Laura K. Chodes, Brooks Applegate, Carol A. Van Hulle, Bennett A. Landman, David H. Zald

## Abstract

Increasing data indicate that prevalent forms of psychopathology can be organized into second-order dimensions based on their correlations, including a general factor of psychopathology that explains the common variance among all disorders and specific second-order externalizing and internalizing factors. Despite this organization, and high levels of comorbidity between diagnoses, most existing studies on the neural correlates of psychopathology employ case-control designs that treat diagnoses as independent categories. Thus, for instance, although perturbations in white matter microstructure have been identified across a range of disorders, the majority of such studies have used case-control designs, leaving it unclear whether observed relations reflect disorder specific characteristics, or transdiagnostic patterns. Using a representative community twin sample of 410 young adults, we tested the hypothesis that some relations between white matter microstructure properties in major tracts are related to second-order factors of psychopathology. We examined fractional anisotropy (FA), radial diffusivity (RD), and axial diffusivity (AD). White matter correlates of all second-order factors were identified after controlling for multiple tests, including the general factor (FA in the body of the corpus callosum), specific internalizing (AD in the fornix), and specific externalizing (AD in the splenium of the corpus callosum, sagittal stratum, anterior corona radiata, and internal capsule). These findings suggest that features of white matter within specific tracts are associated with broad transdiagnostic dimensions of psychopathology rather than being restricted to individual diagnostic categories.

## Introduction

Traditionally, research on psychopathology has focused on specific disorders, and employed case-control designs. This approach has proven problematic given the high degree of heterogeneity within and comorbidity across disorders and the dimensional rather than categorical manner in which psychopathology is expressed (Caspi and Moffitt, 2018; Insel et al., 2010; Lahey et al., 2016). One solution is to characterize psychopathology in terms of latent factors based on the empirically defined organization of symptoms, with second-order factors capturing the transdiagnostic structure of symptoms. Recently, bifactor models have been used as a tool to quantitatively characterize the dimensional structure of psychopathology (Lahey et al., 2015; Lahey et al., 2008). These models include a nonspecific general bifactor on which all prevalent psychiatric disorders load as well as a specific internalizing and specific externalizing factor. The key advantage of this model is that it allows one to disentangle the substantial common variance that is shared across disorders or dimensions (and which has been argued to reflect substantial sharing of etiology across different types of psychopathology), from the variance that is specific to internalizing and externalizing disorders or symptoms (3).

The general factor of psychopathology was initially identified in an adult sample (ages 18-65), but has since been replicated in both children and adolescents (Hankin et al., 2017; Laceulle et al., 2015; Lahey et al., 2015). Caspi and colleagues used data from the Dunedin study to identify a similar model, which includes a general “p-factor” that is defined by shared variance among all disorders (Caspi et al., 2014). When considered at the level of individuals, persons with a broad range of symptoms that cut across second-order dimensions of psychopathology will have a high general factor score, which distinguishes them from persons whose symptoms are limited to just one second-order dimension, such as specific externalizing or specific internalizing symptoms. The extent to which this model of psychopathology proves useful rests on its ability to reveal meaningful features and correlates of psychopathology. In support of this, the general factor predicts both current and future adaptive functioning and demonstrates stability across development (Greene and Eaton, 2017; Lahey et al., 2012; Tackett et al., 2013). Fewer data exist regarding the neural correlates of these dimensions. Such data would be particularly informative because it is difficult to interpret existing case control studies that cannot discriminate between neural correlates that reflect broad shared etiological features or narrower dimensional features of psychopathology. Identifying the neural correlates of shared features of psychopathology will help provide insight into their etiology and may thus yield novel therapeutic targets.

White matter tracts facilitate communication among brain regions, and perturbations in white matter microstructure been consistently identified as a correlate of psychopathology at the disorder level (Thomason and Thompson, 2011). To our knowledge, only one study to date has reported white matter correlates of psychopathology within a bifactor model, and found a relation between white matter microstructure in the cerebellum and the general factor of psychopathology (Romer et al., 2018). While this initial result is promising, there are some notable limitations. For one, they used a college sample at an elite university, and as such these findings might not generalize to either a community sample or one with a wider range of functional impairment. Second, the study employed a whole brain voxel-wise approach rather than a tract specific approach. As such, they may have missed relations at the tract level due to the statistical constraints required for voxel-wise analyses. Finally, they only investigated correlates of the general factor, and did not examine either the specific internalizing or externalizing factors. It is thus unclear if white matter microstructure in specific tracts possesses correlates at the level of either the general factor or the specific internalizing or externalizing factors.

The present study sought to examine whether there are relations between white matter microstructure and second-order factors of psychopathology. We used a representative twin sample with a wide range of psychopathology, examined relations with all factors of psychopathology from the bifactor model, and used a tract specific approach. We hypothesized that given that a number of white matter tracts have been implicated in multiple categorically defined disorders, we would identify relations between a range of white matter tracts and second-order latent factors of psychopathology (Thomason and Thompson, 2011). Given the dearth of studies on this topic to date, we did not formulate hypotheses about specific tracts, and instead examined relations with major white matter tracts of the brain and used false discovery rate corrections to account for the number of tests.

## Methods and Materials

### Participants

Participants were recruited from the Tennessee Twin Study (TTS), which has been conducted in two waves. In the first wave, a representative sample was taken of all live twin births in Tennessee between 1984 and 1995 and consisted of over 2000 twin pairs (Lahey et al., 2008). During this first wave, participants were children and adolescents (ages 8-17) and completed a structured clinical interview along with several other personality measures. During the second wave of the study, twin pairs were young adults (ages 23-31), and were selected with oversampling for internalizing and externalizing psychopathology risk based on the clinical interview from wave one. As such, wave two has a high proportion of individuals with prevalent forms of psychopathology. Individuals were pre-screened and excluded if they had multiple concussions with loss of consciousness or other head injuries, neurological diseases other than headaches, contraindications for MRI scanning, a diagnosis of schizophrenia, or a major developmental disorder. Vanderbilt University’s Institutional Review Board (IRB) approved the study, and the study was conducted in accordance with the guidelines of the IRB including participants providing written informed consent.

A total of 430 young adults in the wave 2 sample completed both a diffusion weighted imaging (DWI) scan and a clinical interview. Twenty participants were excluded for poor DWI data quality (excessive movement, missing data, etc.). The final sample available for analysis consisted of 410 subjects. This included 187 participating twin pairs and 36 individuals whose twin did not provide valid data. There were 92 monozygotic pairs and 95 dizygotic (50 same-sex pairs and 45 different-sex pairs). See Table 1 for demographic characteristics of the sample.

**Table 1.** Participant demographics

| **Variable** | **Mean (Standard Deviation)** |
| --- | --- |
| Age (Years) | 26.05 (1.78) |
| Family income* | 18.79 (4.97) |
| Mother's education (Years) | 13.64 (2.72) |
| **Variable** | **N (Percentage)** |
| Sex |  |
| Male | 196 (47.80) |
| Female | 214 (52.19) |
| Ethnicity |  |
| White | 295 (71.95) |
| African American | 102 (24.88) |
| Other | 13 (3.17) |
| Handedness |  |
| Right | 371 (90.49) |
| Left | 39 (9.51) |
| Scanner |  |
| 3TA | 214 (52.20) |
| 3TB | 196 (47.80) |

\*Family income from wave 1 reported in brackets ranging from 0 (no income) to 24 (\$100,000 and over). 18 = \$35,000 - \$44,999

### Measures

#### Young adult diagnostic interview for children (YA-DISC)

A trained interviewer administered the Young Adult Diagnostic Interview for Children (YA-DISC) to all participants (Shaffer et al., 2000). This computerized structured clinical interview is tailored for the sample’s age range and assesses symptoms from the major diagnostic categories in DSM-IV (Hart et al., 1995; Shaffer et al., 1996). The YA-DISC has the primary advantage that it has few skip-outs, and thus queries symptoms even when the participant cannot reach criteria for a diagnosis, which is critical when measuring dimensional psychopathology.

As implemented in the present study, the YA-DISC assessed symptoms of generalized anxiety disorder (GAD), major depressive disorder (MDD), posttraumatic stress disorder (PTSD), agoraphobia, obsessive-compulsive disorder (OCD), manic episodes, panic attacks, social phobia, specific phobia, as well as nicotine, alcohol, marijuana, and other drug use disorders during the last 12 months. The interview additionally covered disorders that are more frequently associated with adolescent than adult psychopathology, including oppositional defiant disorder (ODD), attention deficit and hyperactivity disorder (ADHD), and conduct disorder (CD).

#### DWI acquisition

Imaging data were acquired on two identical 3T Intera-Achiava Phillips MRI scanners using a 32-channel head coil. T1-weighted images were acquired with a 3-D Magnetization Prepared Rapid Acquisition Gradient Echo (MPRAGE) sequence (TE/TR/TI=4.6/9.0/644(shortest) ms; SENSE=2.0; echo train=131; scan time=4 min 32 s; FOV: 256×256×170 mm, 1 mm isotropic resolution). For DWI, we used a 5 min 2 s multi-slice Stejskal-Tanner spin echo sequence with an echo planar imaging readout (TE/TR=52/7750 ms, SENSE=2.2, FOV: 240×240 mm, 2.5 mm isotropic, 50 slices, 2.5 mm slice thickness). This was acquired with one image without diffusion weighting (“b_0_”) and 32 diffusion-weighted images equally distributed over a hemisphere (b=1000 s/mm^2^).

## Data analysis

### DWI data processing

The DWI data were preprocessed based on methods detailed by Lauzon and colleagues (2013). The fMRIB's Linear Image Registration Tool (FLIRT) from the Functional Magnetic Resonance Imaging of the Brain Software Library (FSL;www.fmrib.ox.ac.uk/fsl) was used to register DWI images to the B0 volume, and then the Brain Extraction Tool (BET) was used to mask the B0 volume (Jenkinson et al., 2002; Smith, 2002). Next FSL was used to perform eddy current and motion corrections. Then the CAMINO software package was used to implement RESTORE robust tensor fitting (Chang et al., 2005; Cook et al., 2006).

After preprocessing the data was quality checked for motion, fractional anisotropy (FA) bias and standard deviation, and goodness of fit of the data to the diffusion model (Lauzon et al., 2013). Subjects were excluded if they were an outlier on any quality assurance metric. Next Tract Based Spatial Statistics (TBSS) were run using FSL, which produced skeletonized white matter images based on the procedures detailed in Smith and colleagues (2006). Subjects’ FA image were first moved to standard space based on a non-linear transformation to the FMRIB58_FA template. Images were then averaged to create a mean FA image, thinned in order to derive a skeletonized mean image, and thresholded at FA > .2. Next, each subject’s FA image was projected onto the mean skeleton, which produced a 4D file that was used for statistical analyses. Finally, both axial diffusivity (AD) and radial diffusivity (RD) skeletonized images were created. This was done by applying the non-linear warp that had been used to bring each FA image to the template, and then applying each subject’s projection vectors onto the mean skeleton. FA measures diffusion broadly whereas AD is more sensitive to properties of axons and RD to property of myelin (Pierpaoli et al., 1996).

We then used the JHU ICBM-DTI white matter labels atlas (Mori et al., 2005) to create masks of the following major white matter tracts: corpus callosum (body, genu, and splenium), corona radiata (anterior, superior, and posterior), internal capsule, external capsule, cingulum, posterior thalamic radiation, uncinate fasciculus, fornix, superior fronto-occipital fasciculus, superior longitudinal fasciculus, and sagittal stratum. Tract masks were overlaid with the white matter skeleton mask generated from the present sample, and only overlapping voxels were included in the final masks. FA, RD, and AD values were averaged from bilateral tracts across all subjects. Values were z-transformed to achieve a mean of 0 and a standard deviation of 1.

### Statistical analyses

All analyses were conducted in Mplus 8.1 and took stratification by age and geographic subareas in sampling procedures and clustering within twin pairs into account in order to adjust for non-independence of twin pairs and sampling methodology (Muthén, 1998-2017). In the first step of all analyses, general, internalizing, and externalizing factors were estimated using a bifactor model with fixed individual disorder dimensions loadings based on the entire wave 2 dataset (n=499) that have been previously published (Lahey et al., 2017b). In the bifactor model all first-order symptom counts were allowed to load onto a general factor. First order symptom counts of inattention, hyperactivity-impulsivity, antisocial personality disorder, mania, and maladaptive nicotine, alcohol, and marijuana misuse all loaded onto the specific externalizing factor and MDD, GAD, PTSD, agoraphobia, panic attacks, social phobia, specific phobia, and mania loaded onto the specific internalizing factor. Because common variance is accounted for by the general bifactor, the specific internalizing and specific externalizing factors are set to be orthogonal. This differs from more traditional correlated factor models in which internalizing and externalizing factor loadings do not distinguish between common and specific sources of variance, and are therefore correlated. The bifactor model included weighting to adjust for both differing probabilities of selection and nonparticipation. Robust maximum likelihood (MLR) estimation was used to account for non-normality in first-order symptom dimensions and adjust standard errors to reflect the clustering of twins within twin pairs.

In the second step, to look at relations between white matter microstructure and latent factors of psychopathology we conducted multiple regressions within structural equation models. Latent factor scores were entered as independent variables, and white matter tract measures (average FA, AD, and RD) across bilateral tracts were dependent variables in separate models. In each model the other latent factors were entered as covariates (e.g. for general factor the specific internalizing and specific externalizing served as covariates). We included the following covariates of no interest: age, sex, ethnicity, scanner and handedness. In order to minimize bias, these analyses applied weights to account for subjects without DWI data, and also accounted for clustering due to the non-independence of twin pairs and stratification based on the age of subjects during the original wave 1 data collection. Significance thresholds were set at *p* < 0.05 using false discovery rate (FDR) within families of tests (FA, AD, and RD) in SPSS in order to account for the large number of tests (45 per family). All raw and processed data are available by request through the RDoC-DB https://data-archive.nimh.nih.gov/rdocdb/.

### Sensitivity analyses

We conducted a series of planned sensitivity analyses to verify the robustness of significant relations. As in the primary analyses, multiple regressions included covariates of no interest, used sampling weights, and accounted for clustering and stratification. We first tested relations separately in males and females given known sex differences in latent factors of psychopathology and in properties of white matter microstructure (Caspi et al., 2014; Hsu et al., 2008). Secondly, we included total intracranial volume (TICV) as a covariate, since some white matter microstructure properties may be impacted by head size (Takao et al., 2011; Takao et al., 2014). TICV was calculated using FreeSurfer (Fischl, 2012). Freesurfer segmentations were visually inspected and edits were made according to the standardized protocols on the software’s website. We excluded data for a total of 8 subjects whose segmentations failed quality assurance checks (excessive movement, processing errors, etc.), and thus analyses with TICV were conducted in a subset of the sample (n = 402).

For the third sensitivity analyses, we tested if findings remained significant with inclusion of the additional demographic covariates of family income and mother’s education from wave 1. In the fourth sensitivity analysis, we looked only at the tracts that showed a significant relation with externalizing, and included a covariate of current drug use (absent or present) as measured on the day of the study visit in a urine-drug test in order to determine if findings were driven solely by substance use. Current substance use was determined based on a urine drug test conducted on the day of testing that included cotinine, amphetamines, methamphetamines, cannabis, methadone, opioids, phencyclidine, barbiturates, benzodiazepines, oxycontin, ecstasy, and propoxyphene. For sensitivity analyses the significance threshold was set to *p* < 0.05 with FDR corrections within families of tests (sex stratified analyses, demographic covariates, TICV, and drug use).

## Results

Participant demographics are presented in Table 1. Multiple regression results are presented in Table 2 including standardized betas, standard errors, and p-values. White matter tracts showing significant relations with latent factors of psychopathology are depicted in Figure 1. The general bifactor had a significant positive relationship with FA in the body of the corpus callosum (CC), such that higher general factor scores are associated with higher FA (*β* = .25, *p* = .001). Microstructure in other tracts did not showed a significant association with the general factor. AD, which scales in the same direction as FA, showed a number of significant negative associations with specific high order factors. The specific externalizing factor was significantly related to AD in the splenium of the corpus callosum (*β* = -.22, *p* = .002), anterior corona radiata (*β* = -.28, *p* = .001), internal capsule (*β* = -.23, *p* = .003), and sagittal stratum (*β* = -.33, p < .001). The specific internalizing factor showed a significant association with AD in the fornix (*β* = -.18, *p* = .003).

**Figure 1.**
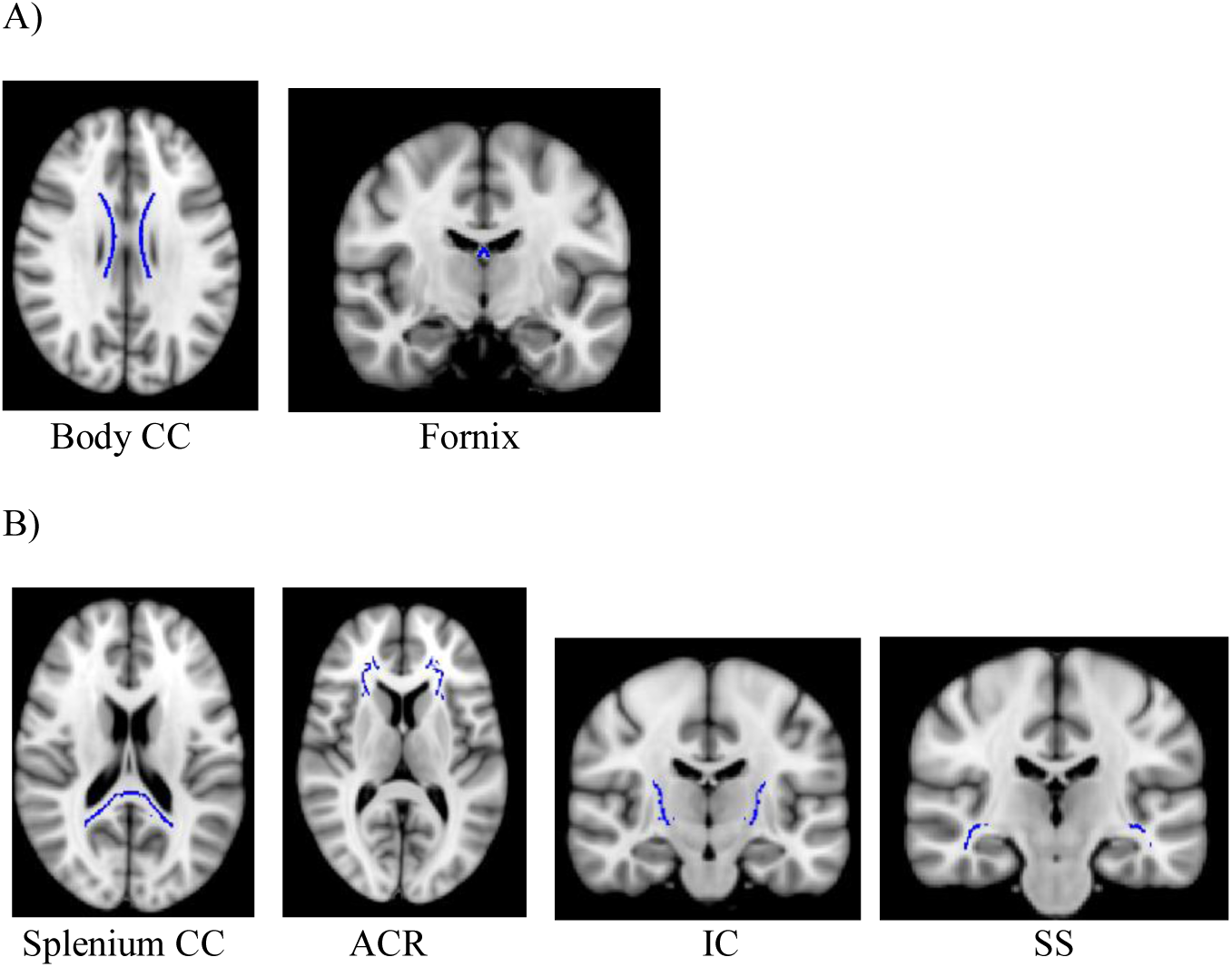
Tracts which showed significant relations with latent factors of psychopathology. A) Left: tract showing significant relation with general factor (body of corpus callosum). Right: tract showing significant relation with specific internalizing (fornix) B) Tracts showing significant relations with the specific externalizing. From left to right: splenium of the corpus callosum, anterior corona radiata, internal capsule, and sagittal stratum.

**Table 2.** Multiple regressions of skeletonized white matter tract indices on latent general and specific internalizing and externalizing factors based on the fixed-loadings bifactor model, controlling demographic covariates of no interest ^a^ (all models n = 410).

| Outcome | Predictor | Fractional anisotropy |  | Radial diffusivity |  | Axial diffusivity |  |
| --- | --- | --- | --- | --- | --- | --- | --- |
|  |  | Regression<br>coeff (SE) | p | Regression<br>coeff (SE) | p | Regression<br>coeff (SE) | p |
| Corpus callosum (body) | **General** | **.25 (0.08)** | **0.001** | -0.25 (0.08) | 0.002 | -0.05 (0.09) | 0.581 |
|  | Internalizing | -0.06 (0.07) | 0.347 | 0.06 (0.07) | 0.424 | -0.03 (0.07) | 0.688 |
|  | Externalizing | 0.08 (0.09) | 0.378 | -0.10 (0.09) | 0.274 | -0.11 (0.10) | 0.238 |
| Corpus callosum (genu) | General | -0.01 (0.06) | 0.823 | -0.04 (0.06) | 0.566 | -0.19 (0.10) | 0.049 |
|  | Internalizing | 0.18 (0.06) | 0.004 | -0.16 (0.07) | 0.014 | 0.04 (0.09) | 0.636 |
|  | Externalizing | 0.01 (0.07) | 0.865 | -0.03 (0.07) | 0.626 | -0.05 (0.10) | 0.581 |
| Corpus callosum (splenium) | General | -0.09 (0.08) | 0.230 | 0.05 (0.08) | 0.480 | -0.11 (0.07) | 0.132 |
|  | Internalizing | 0.04 (0.07) | 0.627 | -0.03 (0.08) | 0.708 | 0.00 (0.07) | 0.949 |
|  | Externalizing | 0.06 (0.08) | 0.437 | -0.15 (0.08) | 0.071 | **-0.22 (0.07)** | **0.002** |
| Anterior corona radiata | General | -0.05 (0.08) | 0.516 | 0.03 (0.08) | 0.742 | -0.09 (0.07) | 0.237 |
|  | Internalizing | 0.12 (0.08) | 0.138 | -0.11 (0.09) | 0.209 | 0.06 (0.08) | 0.435 |
|  | Externalizing | -0.20 (0.09) | 0.019 | 0.04 (0.08) | 0.666 | **-0.28 (0.08)** | **0.001** |
| Superior corona radiata | General | 0.10 (0.10) | 0.293 | -0.08 (0.10) | 0.409 | 0.02 (0.07) | 0.751 |
|  | Internalizing | 0.10 (0.07) | 0.139 | -0.08 (0.07) | 0.213 | 0.05 (0.07) | 0.474 |
|  | Externalizing | 0.03 (0.09) | 0.729 | -0.11 (0.08) | 0.149 | -0.17 (0.09) | 0.055 |
| Posterior corona radiata | General | 0.03 (0.07) | 0.697 | 0.00 (0.06) | 0.974 | 0.02 (0.08) | 0.775 |
|  | Internalizing | 0.12 (0.07) | 0.071 | -0.10 (0.09) | 0.222 | 0.02 (0.09) | 0.839 |
|  | Externalizing | 0.04 (0.10) | 0.674 | -0.11 (0.08) | 0.165 | -0.16 (0.08) | 0.043 |
| Internal capsule | General | -0.02 (0.09) | 0.793 | -0.01 (0.09) | 0.945 | -0.11 (0.08) | 0.132 |
|  | Internalizing | 0.08 (0.07) | 0.283 | -0.10 (0.07) | 0.127 | -0.03 (0.07) | 0.672 |
|  | Externalizing | -0.09 (0.09) | 0.339 | -0.03 (0.10) | 0.727 | **-0.23 (0.08)** | **0.003** |
| External capsule | General | 0.11 (0.10) | 0.253 | -0.11 (0.10) | 0.267 | -0.03 (0.07) | 0.672 |
|  | Internalizing | -0.09 (0.08) | 0.231 | 0.07 (0.10) | 0.488 | -0.07 (0.05) | 0.174 |
|  | Externalizing | -0.04 (0.09) | 0.681 | -0.09 (0.09) | 0.353 | -0.17 (0.07) | 0.016 |
| Cingulum | General | 0.03 (0.10) | 0.750 | -0.02 (0.08) | 0.789 | 0.05 (0.09) | 0.602 |
|  | Internalizing | 0.11 (0.07) | 0.126 | -0.11 (0.07) | 0.077 | -0.02 (0.06) | 0.788 |
|  | Externalizing | -0.12 (0.10) | 0.227 | 0.02 (0.10) | 0.813 | -0.15 (0.09) | 0.095 |
| Posterior thalamic radiation | General | -0.07 (0.08) | 0.388 | 0.07 (0.07) | 0.368 | 0.00 (0.06) | 0.965 |
|  | Internalizing | 0.14 (0.08) | 0.086 | -0.15 (0.08) | 0.072 | -0.02 (0.05) | 0.725 |
|  | Externalizing | -0.03 (0.08) | 0.710 | -0.09 (0.08) | 0.272 | -0.19 (0.07) | 0.010 |
| Uncinate fasciculus | General | 0.05 (0.08) | 0.553 | -0.08 (0.07) | 0.307 | -0.04 (0.08) | 0.673 |
|  | Internalizing | -0.05 (0.06) | 0.388 | 0.06 (0.07) | 0.336 | 0.01 (0.06) | 0.848 |
|  | Externalizing | 0.04 (0.09) | 0.644 | -0.12 (0.09) | 0.185 | -0.09 (0.09) | 0.312 |
| Fornix | General | 0.14 (0.07) | 0.051 | -0.12 (0.07) | 0.069 | -0.07 (0.08) | 0.396 |
|  | Internalizing | 0.12 (0.05) | 0.022 | -0.14 (0.05) | 0.005 | **-0.18 (0.06)** | **0.003** |
|  | Externalizing | -0.08 (0.07) | 0.264 | -0.02 (0.07) | 0.815 | -0.08 (0.07) | 0.249 |
| Superior fronto-occipital fasciculus | General | -0.07 (0.11) | 0.517 | 0.11 (0.09) | 0.200 | 0.03 (0.10) | 0.751 |
|  | Internalizing | -0.02 (0.07) | 0.836 | -0.10 (0.09) | 0.261 | -0.16 (0.08) | 0.046 |
|  | Externalizing | -0.05 (0.12) | 0.697 | -0.04 (0.09) | 0.636 | -0.11 (0.11) | 0.330 |
| Superior longitudinal fasciculus | General | 0.01 (0.08) | 0.867 | -0.02 (0.08) | 0.796 | 0.01 (0.07) | 0.871 |
|  | Internalizing | 0.21 (0.08) | 0.006 | -0.17 (0.07) | 0.024 | 0.12 (0.06) | 0.031 |
|  | Externalizing | -0.02 (0.07) | 0.754 | -0.08 (0.07) | 0.250 | -0.18 (0.09) | 0.034 |
| Sagittal stratum | General | -0.02 (0.09) | 0.777 | 0.03 (0.09) | 0.708 | 0.00 (0.06) | 0.959 |
|  | Internalizing | 0.03 (0.07) | 0.656 | 0.01 (0.08) | 0.949 | 0.03 (0.05) | 0.514 |
|  | Externalizing | -0.14 (0.08) | 0.057 | -0.05 (0.08) | 0.545 | **-0.33 (0.08)** | **0.000** |

<sup>a</sup>Covariates of no interest: Age in wave 2, sex, parent-classified race-ethnicity (Non-Hispanic white versus others), handedness, and scanner; regression coefficients are fully standardized (M = 0, SD = 1). Coefficients in bold were significant after FDR correction within families of 45 tests of significance for each index.

*Sensitivity analyses*

Sensitivity analyses results are presented in Table 3 including standardized betas and standard errors. These analyses confirmed that findings were largely robust to inclusion of additional demographic covariates (mother’s education and family income in wave 1) and were not driven by head size or current substance use (*p*s < .01). In order to test for the presence of interactions with sex, for each of the tracts that showed a significant association in at least one sex, we ran a model in which regression coefficients were allowed to vary by sex for the significant latent factor (e.g. specific internalizing for fornix) and a model in which they were constrained to be equal in the two sexes. We then ran the Satorra-Bentler X^2^ difference test to compare models. These analyses confirmed that there were no significant interactions between sex and psychopathology in their associations with white matter metrics (*p*s > 0.10).

**Table 3.**
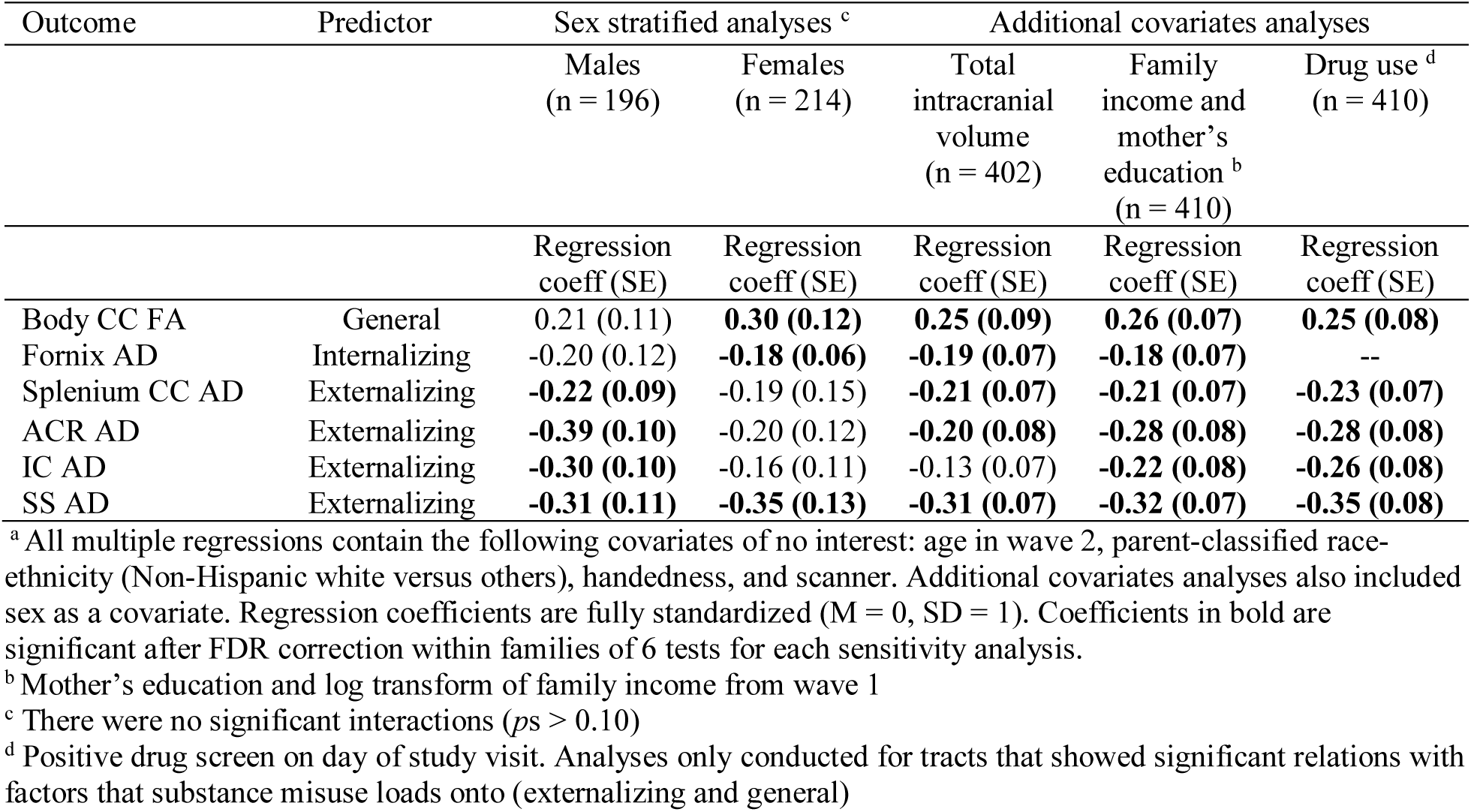
Sensitivity analyses to test robustness of significant relations between white matter tracts and second-order factors of psychopathology. Analyses consist of testing significant relations separately in males and females and inclusion of additional covariates ^a^

| Outcome | Predictor | Sex stratified analyses <sup>c</sup> |  | Additional covariates analyses |  |  |
| --- | --- | --- | --- | --- | --- | --- |
|  |  | Males<br>(n = 196) | Females<br>(n = 214) | Total<br>intracranial<br>volume<br>(n = 402) | Family<br>income and<br>mother's<br>education <sup>b</sup><br>(n = 410) | Drug use <sup>d</sup><br>(n = 410) |
|  |  | Regression<br>coeff (SE) | Regression<br>coeff (SE) | Regression<br>coeff (SE) | Regression<br>coeff (SE) | Regression<br>coeff (SE) |
| Body CC FA | General | 0.21 (0.11) | **0.30 (0.12)** | **0.25 (0.09)** | **0.26 (0.07)** | **0.25 (0.08)** |
| Fornix AD | Internalizing | -0.20 (0.12) | **-0.18 (0.06)** | **-0.19 (0.07)** | **-0.18 (0.07)** | -- |
| Splenium CC AD | Externalizing | **-0.22 (0.09)** | -0.19 (0.15) | **-0.21 (0.07)** | **-0.21 (0.07)** | **-0.23 (0.07)** |
| ACR AD | Externalizing | **-0.39 (0.10)** | -0.20 (0.12) | **-0.20 (0.08)** | **-0.28 (0.08)** | **-0.28 (0.08)** |
| IC AD | Externalizing | **-0.30 (0.10)** | -0.16 (0.11) | -0.13 (0.07) | **-0.22 (0.08)** | **-0.26 (0.08)** |
| SS AD | Externalizing | **-0.31 (0.11)** | **-0.35 (0.13)** | **-0.31 (0.07)** | **-0.32 (0.07)** | **-0.35 (0.08)** |

<sup>a</sup> All multiple regressions contain the following covariates of no interest: age in wave 2, parent-classified race-ethnicity (Non-Hispanic white versus others), handedness, and scanner. Additional covariates analyses also included sex as a covariate. Regression coefficients are fully standardized (M = 0, SD = 1). Coefficients in bold are significant after FDR correction within families of 6 tests for each sensitivity analysis.
<sup>b</sup> Mother's education and log transform of family income from wave 1
<sup>c</sup> There were no significant interactions ( $ps > 0.10$ )
<sup>d</sup> Positive drug screen on day of study visit. Analyses only conducted for tracts that showed significant relations with factors that substance misuse loads onto (externalizing and general)

## Discussion

Applying a bifactor model to characterize psychopathology, we observed relations between second-order factors of psychopathology and several features of white matter microstructure. These data extend a small but growing literature on the neural correlates of latent factor dimensions of psychopathology and demonstrate the potential utility of the general factor model in elucidating biological features that reflect both broad nonspecific aspects of psychopathology as well as those that are specific to a given second-order dimension of psychopathology (Zald and Lahey, 2017). Each latent factor had distinct white matter correlates, providing additional evidence that they are separable at a biological level. These findings join an increasing body of literature indicating associations between transdiagnostic dimensions of psychopathology and other neural measures (Zald and Lahey, 2017).

## Specific externalizing factor correlates

The specific externalizing factor demonstrated the most widespread pattern of correlations, with significant negative associations between this factor and AD in the splenium of the CC, anterior corona radiata (ACR), sagittal stratum (SS), and internal capsule (IC). Decreased integrity in all of these tracts has previously been implicated in case-control studies on antisocial personality disorder (Waller et al., 2017). This parallel is unsurprising given that the antisocial personality symptom dimension loads most highly of all disorders onto the externalizing factor, and as such, antisocial symptomatology is an important aspect of this specific second-order factor (Lahey et al., 2017b). A limitation of prior case-control designs studies is that they did not control for psychopathology outside of the externalizing spectrum. It is thus unclear if identified correlates were specific to externalizing psychopathology or related broadly to non-specific aspects of psychopathology. Based on the present findings, these white matter correlates appear to be specific. Of note, prior studies have identified significant associations with both AD and FA in these tracts, but we only identified relations with AD. Given the typical interpretation of AD, our findings suggest that axonal injury and dysfunction may be especially relevant for dimensional externalizing psychopathology (Song et al., 2003; Song et al., 2002).

The present results also show some concordance with the work of Muetzel and colleagues who observed reduced development in global white matter integrity in relation to externalizing symptoms in childhood (Muetzel et al., 2017). Our findings in young adults may reflect the sequela of an earlier developmental trajectory, although our data suggest that it may be reflected more in specific white matter tracts than a global variable. The specific tracts we identified likely contribute to processes that are similarly affected across externalizing disorders once overlap with internalizing disorders has been covaried.

At present, the mechanisms underlying the link between these specific tracts and externalizing disorders are largely speculative. The splenium is involved in interhemispheric communication, and more specifically connects temporal and occipital areas, and thus is especially relevant for visual processing (Hutchinson et al., 2008; Lansford et al., 2006; Rudge and Warrington, 1991; Schulte and Müller-Oehring, 2010). The SS is implicated more specifically in information processing as it facilitates rapid transfer of information among several cortical regions including the occipital and temporal lobes and subcortical regions including the thalamus (Catani, 2006). At present, it is unclear which specific aspects of visual and information processing these tracts may mediate in the context of externalizing disorders. Some sensory processing weaknesses have been reported in externalizing disorders, especially in terms of processing of facial emotions (Gunn et al., 2009; Marsh and Blair, 2008). Given the regions connected by these tracts, they may also be speculated to contribute to hostile attribution bias, which is a common contributor to aggression in individuals with high externalizing, and has been suggested to involve dysfunction in the way in which visual information is perceived and processed (Bailey and Ostrov, 2008; Lin et al., 2016).

The ACR and IC are both part of the limbic-thalamo-cortical circuitry (Catani et al., 2002; Wakana et al., 2007). The ACR has thalamic projections to regions in the prefrontal cortex that have been implicated in difficulties with top-down emotion regulation control such as the dorsal lateral prefrontal cortex (Karababa et al., 2015; Sanjuan et al., 2013). The IC has projections to the thalamus which in turn is linked to the amygdala, and thus this pathway is likewise relevant for difficulties in top-down emotion regulation (Zikopoulos and Barbas, 2012). Top-down emotion regulation deficits characterize a range of disorders, and as such at present is unclear which aspect is unique to externalizing disorders. It is possible that it is linked to maladaptive emotion regulation in the form of impulsive behavior such as aggression, risk taking, or substance misuse that is observed across externalizing disorders.

Given that problematic substance use is a component of the specific externalizing factor, an important question is the extent to which present findings may represent a consequence of substance use rather than reflect etiology. To address this question, we conducted a sensitivity analysis in which drug use was included as a covariate. All relations remained significant, suggesting that findings were not exclusively driven by substance use. This analysis only accounted for current drug use, and as such further longitudinal study designs are needed to disentangle whether white matter correlates of specific externalizing are a consequence versus a correlate of substance use. Taken together, the present results show reasonable convergence with past studies of different externalizing disorders, but allow a more precise interpretation since the results are not contaminated by more nonspecific features of psychopathology, or impacted by the artificial boundaries of categorical diagnoses.

## Specific internalizing factor correlates

We found a significant negative relation between AD in fornix and the specific internalizing factor. This is consistent with prior studies finding decreased integrity in this tract in MDD and PTSD using case-control designs (Geng et al., 2016; Kennis et al., 2015). As a core pathway of the limbic system, the fornix connects the hippocampus with the mammillary bodies, and is thus believed to be especially important for emotional memory (Aggleton et al., 2005; D’esposito et al., 1995). Impairments in the fornix may contribute to disruptions in emotional memory processes such as fear learning and maladaptive cognitive schema that are expressed across a range of internalizing disorders (Beck, 1983; Britton et al., 2011). That this tract was not also related to the general factor suggests that it is linked more specifically to internalizing psychopathology than to nonspecific aspects of psychopathology.

## General factor correlates

In the present study, we found a significant positive relation between the general factor and FA in the body of the CC. The presence of correlates at the level of the general factor converges with multiple recent neuroimaging and genetics studies that demonstrate that some of the neural correlates of psychopathology reflect broad transdiagnostic aspects rather than being limited to narrower phenotypic features (Goodkind et al., 2015; Kaczkurkin et al., 2017). The CC is critical for interhemispheric communication, and is relevant for multiple cognitive, social and affective processes as it connects a range of regions including precentral areas, parietal and temporal regions, and has projections involving the portions of the insula and cingulate gyrus (Raybaud, 2010; Schulte and Müller-Oehring, 2010).

When considered in the context of most previous studies of the association of variations in CC white matter to mental disorders, the present findings of a positive association of the general factor with CC FA is surprising. Most studies in adult samples that have identified the CC as a correlate of psychopathology have typically identified decreased integrity in patient populations as compared with healthy controls (Arnone et al., 2008; Jiang et al., 2017; Lindner et al., 2016). Nonetheless, some case-control studies of persons given the diagnosis of schizophrenia have found hyperconnectivity between hemispheres (David, 1993; Schmidt et al., 2015). This is important because, in other studies, schizophrenia loads strongly on the general factor of psychopathology (Caspi et al., 2014; Zald and Lahey, 2017).

Notably, some studies of children and adolescents with a broad range of disorders including ADHD, CD, alcohol use disorder, and OCD have also shown a trend similar to the present results with increased white matter integrity in patient populations in the CC, although these results were observed in the context of specific case-control type designs rather than reflecting transdiagnostic characteristics (De Bellis et al., 2008; Decety et al., 2015; Jayarajan et al., 2012; Lawrence et al., 2013; Menks et al., 2017; Pape et al., 2015; Zarei et al., 2011; Zhang et al., 2014). It is notable that in the context of such studies it has been hypothesized that individuals with psychopathology may have an accelerated trajectory of white matter development with an earlier peak and subsequent earlier and steeper decline, thus explaining increased integrity relative to controls during childhood but decreased integrity during adulthood (Menks et al., 2017). The body of the CC is one of the last tracts to complete myelination, not reaching its peak until around age 35 (Lebel et al., 2012). Given that the present sample consists of young adults ages 23-31, it is likely that there is some heterogeneity in the state of development of white matter tracts. Whereas most subjects in this sample have likely reached peak development in all earlier developing tracts, the majority of individuals may not have reached the peak for the body of CC. If the present results reflect a prolonged developmental trajectory in the CC, this might help explain why white matter microstructure properties of the CC body demonstrate findings more in line with pediatric studies. However, the literature on developmental trajectories of white matter and psychopathology shows substantial inconsistencies across studies. For instance, Muetzel and colleagues found that higher early childhood internalizing and externalizing symptoms predicted smaller increases in global FA development later in childhood (Muetzel et al., 2017). By contrast, in a more recent study the same group found there was not a significant association between childhood externalizing symptoms and global FA (Bolhuis et al., 2018). Longitudinal imaging studies will be critical in understanding the complex interplay between white matter tract development trajectories and second-order factors of psychopathology.

## Limitations and conclusions

Based on our scanning parameters we had limited coverage of the cerebellum, and thus were unable to test for a previously reported association between the general factor and white matter microstructure in the cerebellum (Romer et al., 2018). Another limitation is that we excluded individuals who reported psychotic disorders on screening, and therefore more extreme forms of psychopathology were not included in the present sample. Moreover, we did not probe for psychotic symptoms outside of mood congruent symptoms in the context of mood disorders questions. As such, we could not test for correlates of a second-order thought disorder factor, or include thought disorder dimensions in the extraction of the general factor. Studies employing a more comprehensive interview should examine if this putative thought disorder factor has unique white matter microstructure correlates.

The majority of prior studies on white matter microstructure and psychopathology have used case-control designs that obscure the extent to which identified associations reflect nonspecific versus specific aspects of psychopathology. By contrast, the present study applied a bifactor model that allows for a transdiagnostic quantitative approach to distinguish between different second-order dimensions of psychopathology. While there are methodological limitations when using fit statistics to adjudicate between bifactor and traditional correlated factor models of psychopathology, increasing data indicate the ability of this approach to differentiate meaningful nonspecific and dimension-specific correlates of psychopathology including etiological, concurrent and predictive correlates (Caspi and Moffitt, 2018; Lahey et al., 2017a; Wade et al., 2018). Importantly, in the present study, we identified both nonspecific (FA in the body of the CC) and specific (AD in the fornix, sagittal stratum, anterior corona radiata, and splenium of the CC) white matter correlates of psychopathology. Although requiring replication, these results highlight the utility of this quantitative latent factor approach for revealing the neural correlations of psychopathology.

## Acknowledgements

This research was funded by NIMH grant 3R01MH098098-03S1 and Vanderbilt Institute for Clinical and Translational Research (Grant UL1 RR024975-01 & Grant 2 UL1 TR000445-06). This work was supported by the National Science Foundation (NSF) Graduate Research Fellowship Program under Grants Number 0909667 and 1445197 as well as by a National Institute of Mental Health (NIMH) training grant (T32-MH18921). Any opinion, findings, and conclusions or recommendations expressed in this material are those of the authors and do not necessarily reflect the views of the NSF or the NIMH.

